# *In vitro* profiling of orphan G protein coupled receptor (GPCR) constitutive activity

**DOI:** 10.1101/2021.03.10.434788

**Authors:** Lyndsay R. Watkins, Cesare Orlandi

## Abstract

**Background and Purpose:** Members of the G protein coupled receptor (GPCR) family are targeted by a significant fraction of the available FDA-approved drugs. However, the physiological role and pharmacological properties of many GPCRs remain unknown, representing untapped potential in drug design. Of particular interest are ~100 less-studied GPCRs known as orphans because their endogenous ligands are unknown. Intriguingly, disease-causing mutations identified in patients, together with animal studies, have demonstrated that many orphan receptors play crucial physiological roles, and thus, represent attractive drug targets.

**Experimental Approach:** The majority of deorphanized GPCRs demonstrate coupling to G_i/o_, however a limited number of techniques allow the detection of intrinsically small constitutive activity associated with G_i/o_ protein activation which represents a significant barrier in our ability to study orphan GPCR signaling. Using luciferase reporter assays, we effectively detected constitutive G_s_, G_q_, and G_12/13_ protein signaling by unliganded receptors, and introducing various G protein chimeras, we provide a novel, highly-sensitive tool capable of identifying G_i/o_ coupling in unliganded orphan GPCRs.

**Key Results:** Using this approach, we measured the constitutive activity of the entire class C GPCR family that includes 8 orphan receptors, and a subset of 20 prototypical class A GPCR members, including 11 orphans. Excitingly, this approach illuminated the G protein coupling profile of 8 orphan GPCRs (GPR22, GPR137b, GPR88, GPR156, GPR158, GPR179, GPRC5D, and GPRC6A) previously linked to pathophysiological processes.

**Conclusion and Implications:** We provide a new platform that could be utilized in ongoing studies in orphan receptor signaling and deorphanization efforts.

**What is already known:**

- A large group of understudied orphan GPCRs controls a variety of physiological process.

**What this study adds:**

- A new strategy to identify G protein signaling associated with orphan GPCRs.
- Identification of G_i/o_ coupling for 8 orphan GPCRs.

**What is the clinical significance:**

- Many orphan GPCRs are associated with pathological conditions and represent promising druggable targets.

## 1. INTRODUCTION

The large family of G protein coupled receptors (GPCRs) constitutes the most exploited drug target in the human genome (Hauser, Attwood, Rask-Andersen, Schioth, & Gloriam, 2017; Sriram & Insel, 2018). This is the result of GPCR involvement in the regulation of key physiological processes combined with accessibility at the plasma membrane. Nonetheless, the endogenous ligands of many GPCRs have yet to be identified, which collectively are referred to as orphan GPCRs (oGPCRs). In spite of the lack of known endogenous ligands, experimental evidence from both animal models and human studies suggest that many oGPCRs regulate important physiological processes and therefore represent attractive therapeutic targets that remain to be exploited (Audo et al., 2012; Peachey et al., 2012; Wang et al., 2019; Watkins & Orlandi, 2020). The first critical step towards the deorphanization of an oGPCR involves identifying the intracellular signaling pathways that it modulates, thereby providing an essential readout to build screening platforms aimed at testing receptor activation by candidate endogenous/synthetic ligands. However, the lack of known ligands significantly limits the experimental strategies that can be applied to identify oGPCR-activated signaling pathways, thereby representing one of the greatest difficulties in studying oGPCRs. Although not widely utilized, one way to address this question involves measuring the GPCR constitutive activity (Bond & Ijzerman, 2006; Ngo, Coleman, & Smith, 2015). Constitutive activity is observed when a GPCR produces spontaneous G protein activation in the absence of agonist (Rosenbaum, Rasmussen, & Kobilka, 2009), a property often observed when overexpressing GPCRs in heterologous systems and also detected *in vivo* (Corder et al., 2013; Damian et al., 2012; Inoue et al., 2012). Given that current available assays have been unsuccessful in illuminating G protein coupling profiles for many oGPCRs, in particular in detecting those dominantly coupling to G_i/o_ proteins, we sought to develop a novel approach with sufficient sensitivity to study oGPCR pharmacology.

We generated a library of GPCRs that comprises the entire class C GPCR family and a subset of class A members including a total of 19 oGPCRs for testing with several luciferase reporter systems activated in response to G proteins stimulation. Luciferase reporters are characterized by high sensitivity and a wide dynamic range, allowing the detection of even minor levels of G protein-initiated signaling pathways (Cheng et al., 2010). These systems encode either firefly luciferase or nanoluc under the control of inducible promoters downstream of the main G protein-promoted signaling cascades. Gα proteins are classified into four major families: G_s_, G_q_, G_12/13_, and G_i/o_. In detail, activation of G_s_ family members (G_s/olf_ stimulates adenylate cyclase to produce cAMP that triggers downstream signaling events which activate the cAMP response element (CRE). A primary effector of heterotrimeric G_q_ family members (G_q/11/14/15_) is phospholipase Cβ (PLCβ) that catalyzes the formation of second messengers inositol 1,4,5-trisphosphate and diacylglycerol leading to the activation of the Nuclear Factor of Activated T-cells (NFAT) promoter. The canonical downstream target of the heterotrimeric G_12/13_ proteins is a group of Rho guanine nucleotide exchange factors (RhoGEFs) that activate the Ras-family small GTPase RhoA. G_12/13_ activation can be detected by luciferase reporters using promoters comprising a serum response element (SRE), or serum response factor response element (SRF-RE), with the last one designed to respond to SRF-dependent and ternary complex factor (TCF)-independent pathways (Cheng et al., 2010). Conversely, detection of active G proteins belonging to the G_i/o_ family (G_i1/i2/i3/o/z/t_) is more complicated and elusive. In fact, the main effect of G_i/o_ stimulation consists in the inhibition of adenylate cyclase, leading to a reduction of the cAMP production. Changes in cAMP levels can be readily detected after agonist-stimulation of G_i/o_-coupled GPCRs, however, determining the constitutive activity of such receptors has proven challenging. Moreover, according to the GPCR database (Flock et al., 2017; Pandy-Szekeres et al., 2018) (https://gpcrdb.org/signprot/statistics_venn), 158 out of 247 ligand-activated GPCRs (64%) can activate members of the G_i/o_ protein family, with half of them, 79 out of 247 (32%), showing exclusive coupling to G_i/o_. Considering the likely large number of G_i/o_-coupled receptors among oGPCRs, research in this field is in desperate need of innovative sensitive tools.

To overcome this issue, we took advantage of previously developed G protein chimeras (Ballister, Rodgers, Martial, & Lucas, 2018; Conklin, Farfel, Lustig, Julius, & Bourne, 1993; Inoue et al., 2019), and generated novel G protein chimeras to expand the GPCR toolkit. Swapping the C-terminal strand of amino acids of any G protein with those of G_i/o_ family members enables GPCRs that preferentially couple to G_i/o_ to trigger alternative downstream signaling events (Ahmad, Wojciech, & Jockers, 2015; Ballister et al., 2018; Conklin et al., 1993; Coward, Chan, Wada, Humphries, & Conklin, 1999; Inoue et al., 2019). Exploiting such property, we rerouted pathways initiated by the constitutive activation of G_i/o_-coupled receptors to different downstream signaling outcomes which are more readily measurable. Herein, we tested 8 G protein chimeras against a set of 8 well-characterized G_i/o_-coupled receptors for a total of 64 combinations to identify the most suitable ones for analysis of constitutive activity across our oGPCR library. Applying this strategy we successfully identified 8 oGPCRs that show significant basal activation of G_i/o_ proteins. We finally validated these results by measuring the inhibition of forskolin-induced cAMP production by an oGPCR showing high G_i/o_ constitutive activity, GPR156.

## 2. METHODS

### 2.1 Cell cultures and transfections

HEK293T/17 cells were cultured at 37°C and 5% CO_2_ in Dulbecco’s Modified Eagle’s Medium (DMEM; Gibco, 10567-014) supplemented with 10% fetal bovine serum (FBS; Biowest, S1520), Minimum Eagle’s Medium (MEM) non-essential amino acids (Gibco, 11140-050), and antibiotics (100 units/ml penicillin and 100 μg/ml streptomycin; Gibco, 15140-122). HEK293 cells were seeded in 6-well plates in medium without antibiotics at a density of 1 × 10^6^ cells/well. After 4 hours, cells were transfected using linear 25 kDa polyethylenimine (PEI) (VWR; AAA43896) at a 1:3 ratio between total μg of DNA plasmid (2.5 μg) and μl of PEI (7.5 μl). A pcDNA3.1 empty vector was used to normalize the amount of transfected DNA. For western blot and BRET assays, cells were collected 24 hours after transfection. For CRE and NFAT luciferase reporter assays, cells were incubated overnight and then serum-starved in Opti-MEM Reduced Serum Media (Gibco, 11058-021) for 4 hours before collection. For SRE and SRF-RE luciferase reporter assays, cells were incubated overnight and then serum-starved in Opti-MEM for 24 hours before collection.

### 2.2 DNA constructs and cloning

Details about the DNA constructs used in this paper are listed in the Supplementary table 1. Plasmids encoding GPR158, GPR179, ADRA2A, LPAR2, CHRM1, GRM1, GRM2, GRM3, GRM4, GRM6, GRM7, GRM8, GABBR1, GABBR2, masGRK3CT-Nluc, Gα_i1_, Gα_i3_, Gα_oA_, and Gα_z_ were generous gifts from Dr. Kirill Martemyanov (The Scripps Research Institute, FL). The plasmid encoding the human GRM5a was a kind gift from Dr. Paul Kammermeier (University of Rochester, NY). Gβ1-Venus156-239 and Gγ2-Venus1-155 were generous gifts from Dr. Nevin Lambert (Augusta University, GA) (Hollins, Kuravi, Digby, & Lambert, 2009). Plasmids encoding the cAMP sensor (pGloSensor-22F) and the following luciferase reporters were purchased from Promega: CRE-luc2, CRE-Nluc, NFAT-luc2, NFAT-Nluc, SRE-luc2, and SRF-RE-luc2. The plasmid encoding for the renilla luciferase under control of the constitutively active thymidine kinase promoter (pRL-tk) was a kind gift from Dr. Mark Ginsberg (University of California San Diego, CA). Plasmids encoding the following GPCRs were obtained from cDNA Resource Center (www.cdna.org): ADRB2, HTR1A, HTR2A, HTR4, and DRD1. The following cDNA clones from the Mammalian Gene Collection (MGC) encoding for full-length GPCR sequences required to further subcloning were purchased from Horizon Discovery: GPR19, GPR37, GPR85, GPR137, GPR137b, GPR162, GPR176, GPR180, CaSR, GPR156, GPRC5A, GPRC5B, GPRC5C, and GPRC6A. Codon optimized sequences for the following oGPCRs used to further subcloning were a kind gift from Dr. Bryan Roth (University of North Carolina, NC) (Kroeze et al., 2015): GPR22 (Addgene plasmid #66346), GPR88 (Addgene plasmid #66380), GPR151 (Addgene plasmid #66327). The plasmids encoding the following G_q_-derived chimeras were a kind gift from Dr. Bruce Conklin (University of California San Francisco, CA) (Conklin et al., 1993): qo5 (Addgene plasmid #24500), qi_1_5 (Addgene plasmid #24501), qz5 (Addgene plasmid #25867). The plasmids encoding the following G_s_-derived chimeras were a kind gift from Dr. Robert Lucas (University of Manchester, UK) (Ballister et al., 2018): Gsz (Addgene plasmid #109355), Gso (Cys) (Addgene plasmid #109375), Gsi (Cys) (Addgene plasmid #109373). The plasmids encoding the following GPCRs were a kind gift from Dr. Erik Procko (University of Illinois at Urbana, IL) (Park et al., 2019): HLA-cMyc-EcopT1R1 (Addgene plasmid #113962), HLA-Flag-natT1R3 (Addgene plasmid #113950), HA-Flag-natT1R2 (Addgene plasmid #113944). The codon optimized sequence for human GPRC5D expression in mammalian cells was synthetized by Integrated DNA Technologies as a gene block and inserted into a pcDNA3.1 vector including a C-terminal HA-tag using In-Fusion HD Cloning technology (Clontech). The full-length sequences of all the orphan GPCRs (except GPR158 and GPR179) were subcloned into a pcDNA3.1 vector for mammalian expression and a C-terminal HA-tag (YPYDVPDYA) was add using In-Fusion HD Cloning technology (Clontech). A plasmid encoding the G protein chimera G_q_G_i3_ bearing the core of human Gαq and the last 4 amino acid of Gα_i3_ was generated by primer mutagenesis and In-Fusion HD Cloning (Clontech) in a pcDNA3.1 vector. All constructs were verified by Sanger sequencing.

### 2.3 Western blot

For Western blotting analysis, transfected cells were harvested and lysed by sonication in icecold immunoprecipitation buffer (300 mM NaCl, 50 mM Tris-HCl, pH 7.4, 1% Triton X-100, and complete protease inhibitor mixture). Lysates were cleared by centrifugation at 14,000 rpm for 15 min, and the supernatants were diluted in SDS sample buffer (final concentrations: 50 mM Tris-HCl pH 6.8, 1% SDS, 10% glycerol, 143 mM 2-mercaptoethanol, and 0.08 mg/ml bromophenol blue). 10 μl of each protein sample were loaded and analyzed by SDS-PAGE. Orphan GPCR expression was detected using rat anti-HA tag (clone 3F10) antibodies (Sigma-Aldrich; 11867423001) or rabbit anti-myc tag antibodies (GenScript; A00172).

### 2.4 Luciferase reporter assays

HEK293T/17 cells were plated at a density of 1 × 10^6^ cells/well in 6-well plates in antibiotic-free medium and transfected as described above. 2.5 μg of total DNA plasmids were transfected according to the following ratio: 0.97 μg of pRL-tk plasmid expressing renilla luciferase under control of the constitutive thymidine kinase promoter; 0.14 μg of luciferase reporter (NFAT-Fluc, CRE-Fluc, SRE-Fluc, and SRF-RE-Fluc for screening of G_q_, G_s_, and G_12/13_ activation; NFAT-Nluc and CRE-Nluc for screening of G_i/o_ activation); 1.11 μg of GPCR; and only in experiments screening G_i/o_ activation, 0.28 μg of G protein chimeras (G_q_G_i1_, G_q_G_i1_-9, G_q_G_i3_, G_q_G_o_, G_q_G_z_, G_s_G_i1_, G_s_G_o_, or G_s_G_z_). pcDNA3.1 was used to normalize the amount of transfected DNA. For CRE and NFAT luciferase reporter assays, cells were incubated overnight and then serum-starved in Opti-MEM for 4 hours before collection. For SRE and SRF-RE luciferase reporter assays, cells were incubated overnight and then serum-starved in Opti-MEM for 24 hours before collection. Transfected cells were harvested, centrifuged for 5 minutes at 500g, and resuspended in 500 μl of PBS containing 0.5 mM MgCl_2_ and 0.1% glucose. 50 μl of cells were incubated in 96-well flat-bottomed white microplates (Greiner Bio-One) with 50 μl of luciferase substrate according to manufacturers’ instructions: furimazine (Promega NanoGlo; N1120) for nanoluc, e-coelenterazine (Nanolight; 355) for renilla luciferase, and luciferin (Promega BrightGlo; E2610) for firefly luciferase. Luciferase levels were quantified using a POLARstar Omega microplate reader (BMG Labtech). Renilla luciferase expression was used to normalize the signal in order to compensate for variability due to transfection efficiency and number of cells.

### 2.5 Bioluminescence Resonance Energy Transfer (BRET) assays

Measurements of ADRA2A activation by norepinephrine in live cells by measurement of BRET between Venus-Gβ1γ2 and masGRK3CT-Nluc was performed as described previously (Masuho, Martemyanov, & Lambert, 2015). 2.5 μg of total DNA plasmids were transfected according to the following ratio: 0.21 μg of Gβ1-Venus156-239; 0.21 μg of Gγ2-Venus1-155; 0.21 μg of masGRK3CT-Nluc; 0.42 μg of Gα_i/o_ proteins or G_q_-derived chimeras (Gα_i1_, Gα_i3_, Gα_o_, Gα_z_, G_q_G_i1_, G_q_G_i1_-9, G_q_G_i3_, G_q_G_o_, G_q_G_z_) or 1.25 μg of Gα_s_-derived chimeras (G_s_G_i_, G_s_G_o_, or G_s_G_z_); and 0.21 μg of ADRA2A. Empty vector pcDNA3.1 was used to normalize the amount of transfected DNA. 18 hours after transfection, HEK293T cells were washed once with phosphate-buffered saline (PBS). Cells were then mechanically harvested using a gentle stream of PBS, centrifuged at 500 g for 5 minutes, and resuspended in 500 μl of PBS containing 0.5 mM MgCl_2_ and 0.1% glucose. 25 μl of resuspend cells were distributed in 96-well flat-bottomed white microplates (Greiner Bio-One). The nanoluc substrate furimazine (N1120) was purchased from Promega and used according to the manufacturer’s instructions. BRET measurements were obtained using a POLARstar Omega microplate reader (BMG Labtech) which permits detection of two emissions simultaneously with the highest possible resolution of 20 ms per data point. All measurements were performed at room temperature. The BRET signal was determined by calculating the ratio of the light emitted by Venus-Gβ1γ2 (collected using the emission filter 535/30) to the light emitted by masGRK3CT-Nluc (475/30). The average baseline value (basal BRET ratio) recorded for 5 seconds before agonist application was subtracted from the BRET signal to obtain the ΔBRET ratio.

### 2.6 cAMP assay

HEK293T cells were transfected with an equal ratio of indicated GPCR plasmid and pGloSensor™-22F cAMP plasmid (Promega). 18 hours post-transfection, cells were detached with 1 ml of PBS, centrifuged at 500 g for 5 minutes, and resuspended in 300 μl of PBS containing 0.5 mM MgCl_2_ and 0.1% glucose. 40 μl of the cell suspension were transferred to each well of 96-well plates containing 10 μl of 5X GloSensor cAMP Reagent (Promega) prepared according to the manufacturer’s instruction. Cells were then incubated at 37°C for 2 hours and let cool down to room temperature for 10 minutes. Luminescence was monitored every 30 seconds using a POLARstar Omega microplate reader (BMG Labtech) at room temperature. After 3 minutes, forskolin (Tocris; 1099) was added to the cells at a final concentration of 0.5 μM.

### 2.7 Statistical analysis

Analyses were performed using GraphPad Prism 9 software and number of biological and technical replicates are described in the figure legends. Data in figure 4 were analyzed by normalizing the nanoluc/renilla luciferase ratio by control cells not transfected with the G protein chimeras. One-way ANOVA with Dunnett’s multiple comparisons test was performed comparing the signal obtained with each oGPCRs against control cells not expressing GPCRs.

## 3. RESULTS

### 3.1 Screening of GPCR constitutive activity using luciferase reporter assays

Taking advantage of a plasmid cDNA library encoding the entire 22 class C GPCRs, including 8 oGPCRs, and a subset of 19 class A GPCRs, including 11 oGPCRs, we systematically tested their constitutive activity using several luciferase reporter assays and setting an arbitrary threshold of 3 fold-increase for positive signals (Figure 1). We first screened the library for G_q_ activation by co-transfecting HEK293T cells with each GPCR and a NFAT-RE luciferase reporter (Figure 1a). As expected, the positive controls serotonin 2A receptor (HTR2A), muscarinic receptor 1 (CHRM1), and the metabotropic glutamate receptors 1 and 5 (GRM1 and GRM5) showed the highest signal compared to control cells expressing only the luciferase reporter. None of the oGPCRs tested showed any constitutive activation of G_q_-dependent signaling pathways (Figure 1a). Similarly, using a CRE reporter assay for G_s_ activation, we detected the constitutive activity of β2 adrenergic receptor (ADRB2), serotonin receptor 4 (HTR4), and dopamine receptor D1 (DRD1) (Figure 1b). Interestingly, some G_q_-coupled receptors also triggered the expression of luciferase activating the CRE reporter, while all the oGPCRs showed levels of activation comparable to those of the control not expressing GPCRs (Figure 1b). Using the same approach with two additional luciferase reporters SRE and SRF-RE, we detected the constitutive activity of the lysophosphatidic acid receptor 2 (LPAR2) downstream of G_12/13_ (Figure 1c-d). Again, no constitutive luciferase expression by oGPCRs was detected (Figure 1c-d). Overall, these experiments revealed that luciferase reporters are sensitive enough to detect GPCR activity in the absence of ligand application for some GPCRs; however, none of the 19 oGPCRs tested showed any constitutive activation of G_q_, G_s_, or G_12/13_ signaling pathways.

**Figure 1.**
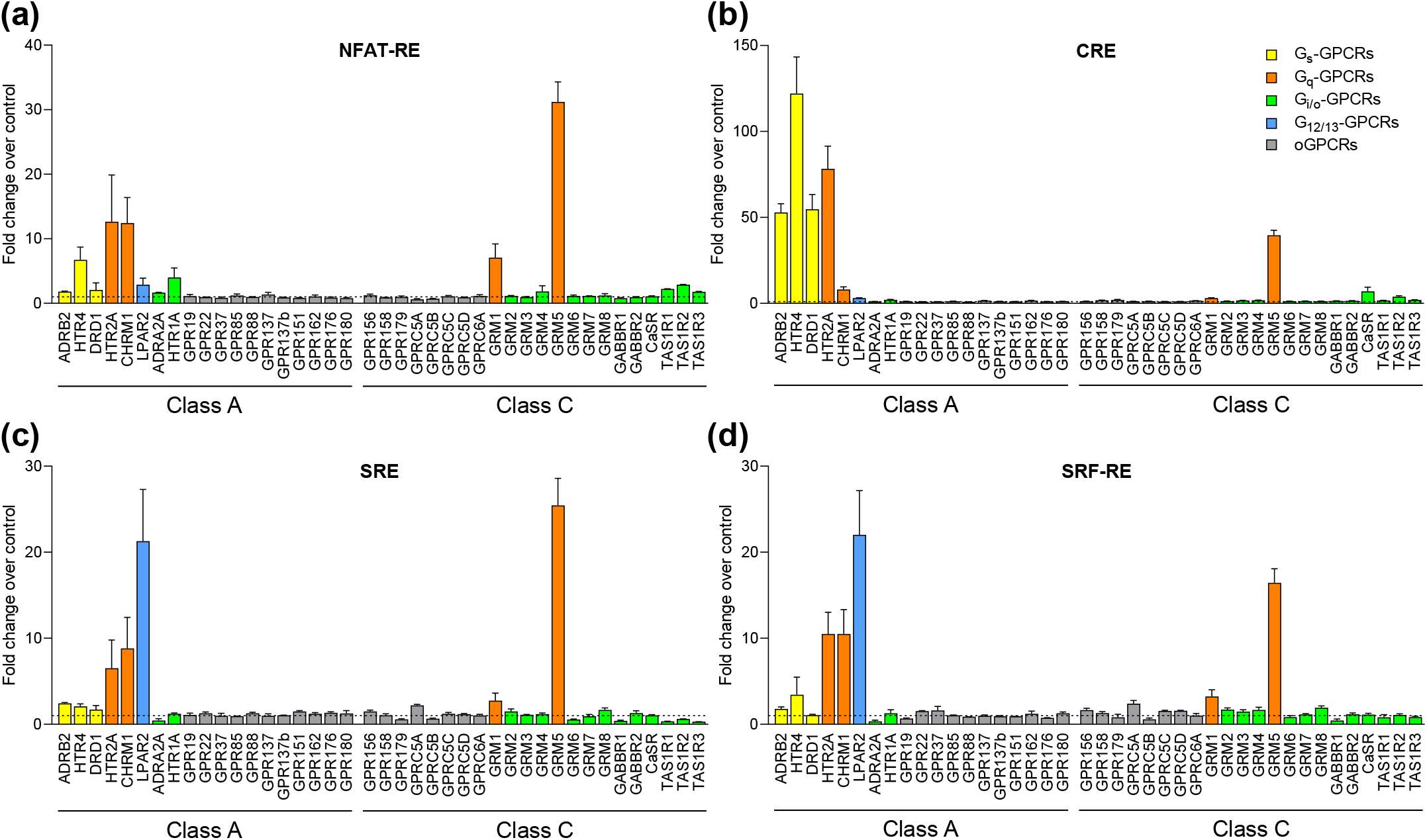
Analysis of GPCR constitutive activity. 4 luciferase reporter assays were used to measure the constitutive activity of a library of 41 GPCRs that include 19 orphan GPCRs. The amount of firefly luciferase accumulated in the cells was normalized on the levels of constitutively expressed renilla luciferase. The effect of GPCR overexpression was compared to cells expressing only the reporters (control cells, dotted line) and reported as fold-change over control. Colors are used to discriminate know G protein coupling: G_s_ (yellow), G_q_ (orange), G_12/13_ (blue), and G_i/o_ (green). oGPCRs are in gray. (a) NFAT-RE-induced luciferase expression. (b) CRE-induced luciferase expression. (c) SRE-induced luciferase expression. (d) SRF-RE-induced luciferase expression. The data shown represent the average of 3-6 independent experiments, each performed in duplicate. Data shown as means ± SEM.

### 3.2 G protein chimeras are valuable tools to detect constitutive activity of G_i/o_-coupled receptors

After exploring our GPCR library for the activation of signaling pathways downstream of G_s_, G_q_, and G_12/13_, we focused on G_i/o_ signaling. In principle, the CRE luciferase reporter could be used to detect activation of G_i/o_-coupled GPCRs as a reduction in cAMP levels, however its use is limited by a low dynamic range. This especially applies to measurements of GPCR constitutive activity, as they are intrinsically small. Therefore, to obtain a reliable quantification of constitutive activation of G_i/o_ signaling, we tested several G protein chimeras based on G_q_ or G_s_ core protein and bearing the C-terminus of either G_i1_, G_i3_, G_o_, or G_z_ (Figure 2a). The last few amino acids in the Gα protein C-terminus define most of the GPCR coupling selectivity (Ballister et al., 2018; Conklin et al., 1993; Inoue et al., 2019). However, the coupling efficiency of G protein chimeras is variable and depends on the GPCR analyzed (Ballister et al., 2018; Conklin et al., 1993; Inoue et al., 2019). Thus, we tested 5 chimeras based on a G_q_ core and 3 chimeras based on a G_s_ core for their ability to stimulate NFAT or CRE luciferase reporters, respectively. As a control, we first quantified the amount of luciferase expressed in cells where each chimera was co-transfected with the associated luciferase reporter but without GPCR overexpression. We reasoned that the difference in luciferase expression obtained comparing cells expressing the reporter with or without expression of the G protein chimeras could represent an index of reporter activation by endogenously expressed G_i/o_-coupled receptors. As a positive control, we co-transfected GRM2 because of its reported high constitutive activity (Doornbos et al., 2018) (Figure 2b-c). We found that expression of G_q_-based chimeras only produced a negligible amount of NFAT reporter induction (0.1-fold increase on average) (Figure 2b), while we observed an average of 83-fold increase in CRE-induced luciferase expression using the G_s_-based chimeras (Figure 2c). As expected, expression of GRM2 significantly induced the luciferase expression with all of the chimeras tested (Figure 2b-c). The fold-change observed normalizing the GRM2 constitutive activity over the no-GPCR control, revealed comparable levels of activation between G_q_ and G_s_ chimeras. Interestingly, the G_q_G_i3_ chimera showed a 29.3 ± 3.5 fold-increase, being the highest amplitude among all the chimeras, while both the G_q_G_z_ and the G_s_G_z_ chimeras showed only a 4.7 ± 0.6 and 3.8 ± 0.4 fold-increase (Figure 2b-c). To explore the efficiency of activation of these G protein chimeras, we then quantified the constitutive activity of eight GPCRs that are known to primarily couple to G_i/o_ (Flock et al., 2017; Pandy-Szekeres et al., 2018): class C GRM2, GRM3, GRM4, GRM6, GRM7, GRM8, and class A ADRA2A and HTR1A (Figure 2d). Assuming an arbitrary threshold of 3 fold-increase as a positive signal, our data show that some GPCR constitutive activity can be detected with the majority of the chimeras (i.e. GRM2), while some GPCRs show levels of activation above the threshold only if co-transfected with G_s_G_i1_ or G_s_G_o_ chimeras (i.e. GRM7 and GRM8). According to earlier reports, the signal amplitude is GPCR-dependent (Conklin et al., 1993), and here we show it is undetectable for a subset of GPCR (i.e. ADRA2A). We next asked if the absence of signal for ADRA2A could be due to a lack of constitutive activity or to the expression of a non-functional receptor. Using a Bioluminescence Resonance Energy Transfer (BRET) assay we tested the ligand activation of the G protein chimeras by ADRA2A (Figure 3a). Here, we activated the ADRA2A receptor with the endogenous agonist norepinephrine at a concentration of 1 μM. We compared the ΔBRET ratio obtained using G protein chimeras with those obtained with wild type G_i1_, G_i3_, G_o_, and G_z_ proteins (Figure 3b-d). Although the amplitude of the BRET signal generated by the G protein chimeras was smaller compared to the signal produced by wild type G protein, our data show that ADRA2A can indeed activate every tested chimera. Overall, we provide evidence that ADRA2A lacks detectable levels of constitutive activity. Likewise, we expect that the constitutive activity of some of the oGPCRs in our library will also be undetectable.

**Figure 2.**
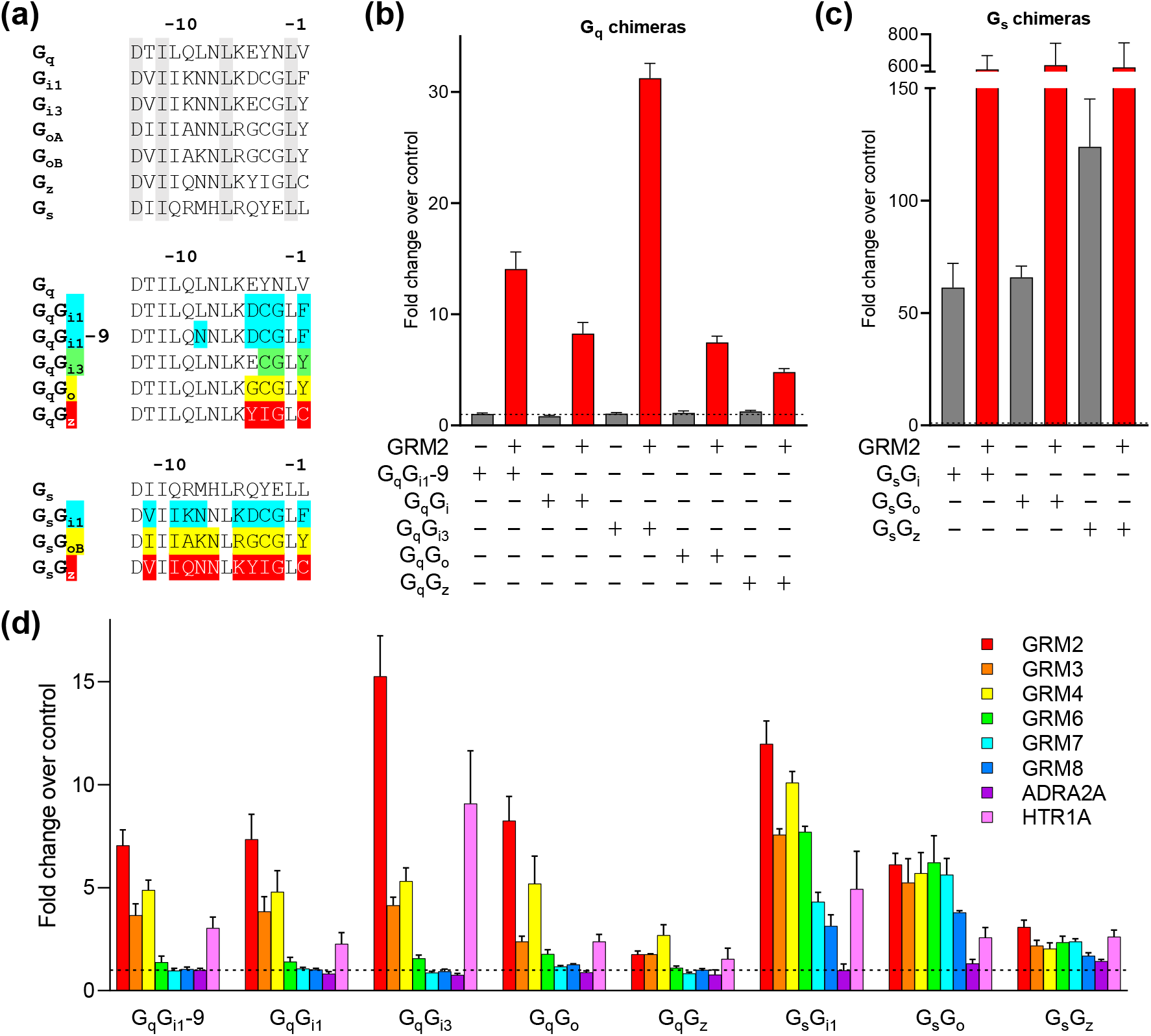
Use of G protein chimeras to detect constitutive G_i/o_ activation. (a) Sequence alignment of the last 14 amino acids of wild type G proteins (top), G_q_-derived chimeras (middle), and G_s_-derived chimeras (bottom). Residues conserved among every G protein are highlighted in gray. Residues that were substituted in the G protein chimeras were highlighted and aligned with the G protein C-terminal sequence of the core protein (G_q_, middle; G_s_, bottom). (b) Constitutive activation of G_q_ chimeras by overexpression of GRM2. Reported is the fold change over control cells expressing only NFAT-Nluc reporter and renilla luciferase (dotted line). (c) Constitutive activation of G_s_ chimeras activate by GRM2 and reported as fold change over control cells expressing only CRE-Nluc reporter and renilla luciferase (dotted line). (d) Analysis of the constitutive activity of the G_i/o_-coupled metabotropic glutamate receptors GRM2, GRM3, GRM4, GRM6, GRM7, and GRM8, the α2A-adrenergic receptor (ADRA2A), and the serotonin 1A receptor (HTR1A) with each of the 8 G_q_-and G_s_-based chimeras. The data shown represent the average of 3 (panels b and c) or 3-6 (panel d) independent experiments, each performed in duplicate. Data shown as means ± SEM.

**Figure 3.**
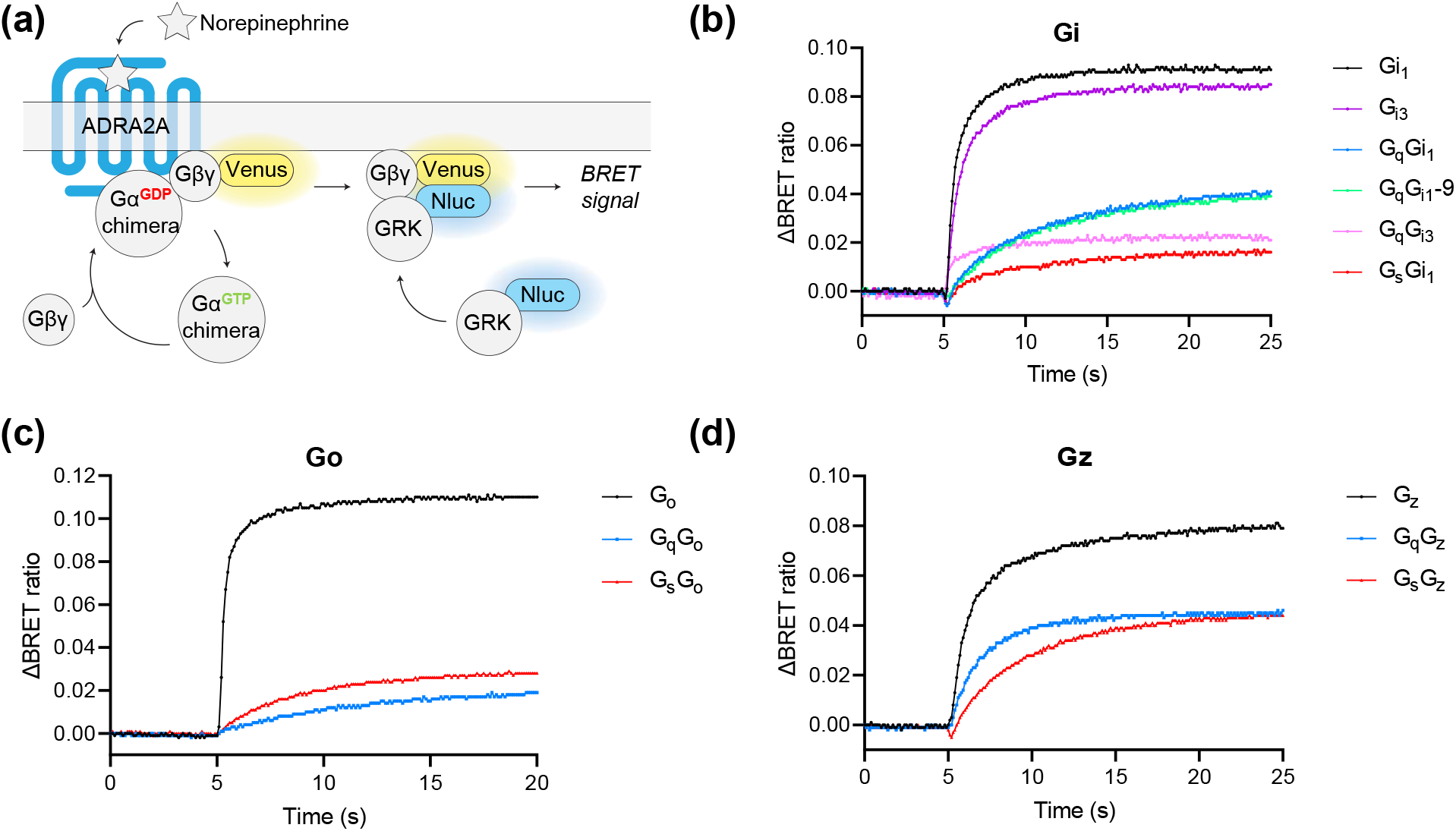
G protein chimera activation by agonist-activated GPCRs using BRET assay. (a) Schematic representation of the BRET assay used to detect agonist-induced activation of ADRA2A. Norepinephrine application triggers the GDP exchange with GTP on the Gα subunit and the subsequent dissociation of Gβγ-Venus. At the membrane, released GßY will interact with the C-terminus of masGRK3 that is fused with Nanoluc. Using a BMG Omega plate reader we can therefore detect the BRET signal generated. (b) Representative response profile showing the BRET signal after norepinephrine application at 5 seconds. The ΔBRET ratio is calculated for each of the wild type G_i1_ and G_i3_, or the chimeric G proteins bearing a G_i1_ or G_i3_ C-terminus. (c) Norepinephrine activation of wild type G_o_ or G protein chimeras with a G_o_ C-terminus. (d) Norepinephrine activation of wild type G_z_ or G protein chimeras with a G_z_ C-terminus. The data shown were replicated in 3 independent experiments.

**Figure 4.**
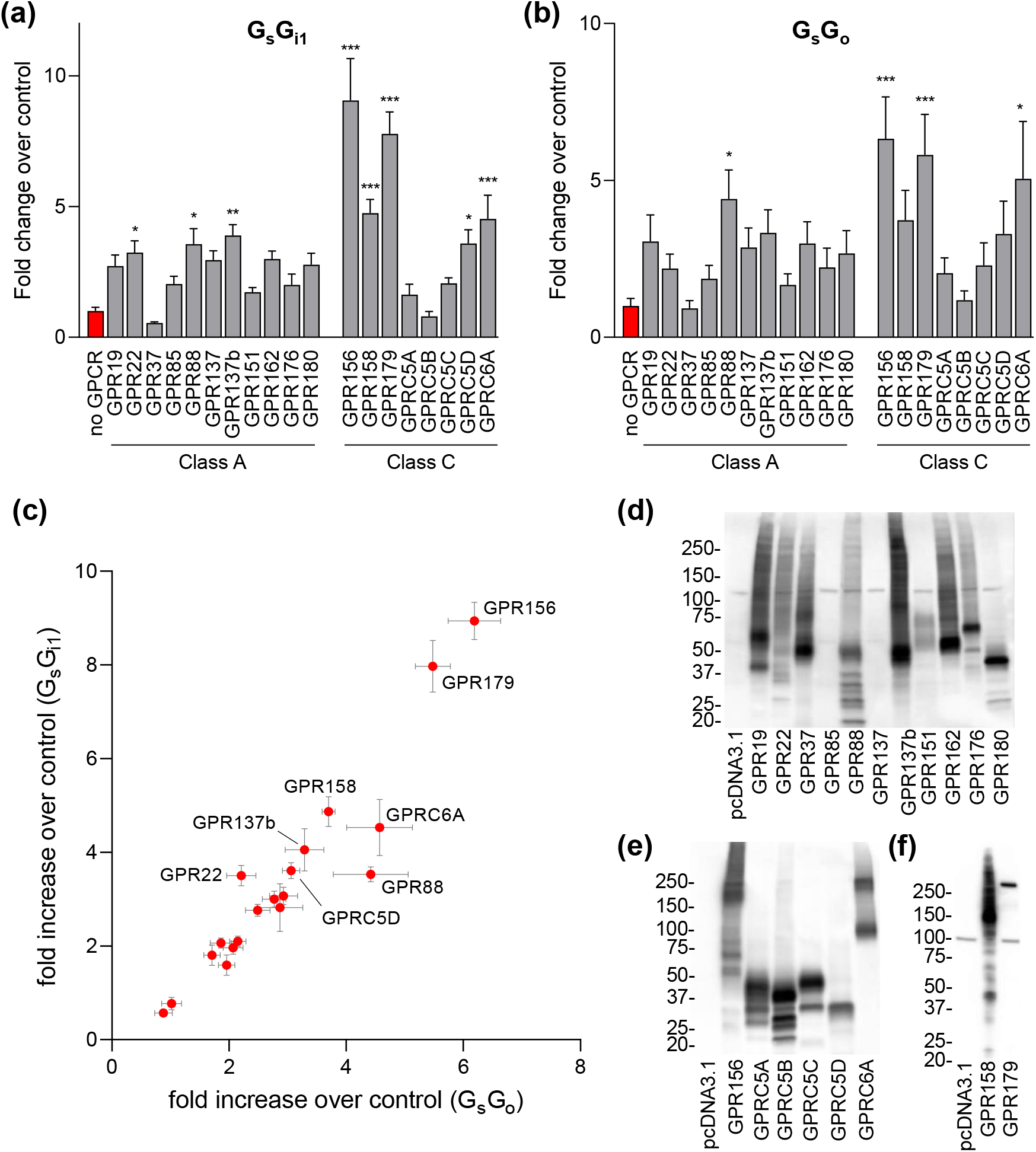
Orphan GPCR screening for G_i/o_ coupling. Quantification of luciferase expression in cells co-transfected with oGPCRs, G_s_-based chimeras, CRE-Nluc, and renilla luciferase showed as fold change over control cells not overexpressing oGPCRs. (a) Analysis of constitutive activation of the G_s_G_i1_ chimera by class A and class C oGPCRs. (b) Constitutive activation of the G_s_G_o_ chimera by class A and class C oGPCRs. (c) Bi-dimensional representation of oGPCR G_i/o_ constitutive activity. Only oGPCRs showing statistically significant activity are labeled. The data shown represent the average of 4 (G_s_G_i1_) or 5 (G_s_G_o_) independent experiments, each performed in duplicate (one-way ANOVA with Dunnett’s multiple comparisons test, *p<0.05, **p<0.01 ***p<0.001). Data are shown as means ± SEM. (d-f) Western blot analysis of protein levels detected in cells transfected with oGPCRS or without (pcDNA3.1) as a negative control. Antibodies raised against HA tag were used to detect class A oGPCRs (d), and some class C oGPCRs (e). Antibodies against myc tag were used to detect class C GPR158 and GPR179 (f).

### 3.3 Identification of oGPCRs that signal through G_i/o_

Agonist-activation of a subset of G_i/o_ coupled receptors, M4R, D2R, α2AAR, and A1R, using G protein chimeras was previously reported to be strongly dependent on both the G_i/o_ protein core and the G_i/o_ C-terminus (Okashah et al., 2019). However, among the possible combinations of G protein cores and C-termini, it was established that chimeras based on G_s_ could be triggered by G_i/o_ coupled receptors more easily than chimeras bearing the core of G_q_ or G_12/13_ (Okashah et al., 2019). Our data on 8 control GPCRs suggest similar preference pattern, with G_s_ chimeras being more promiscuous than G_q_ chimeras (Figure 2d). We thus screened our oGPCR library for constitutive activation of G_s_G_i1_ and G_s_G_o_ chimeras normalizing the luciferase signal to that obtained in cells transfected only with the luciferase reporters but no G protein chimeras (Figure 4a-b). Excitingly, this optimized assay indicated that 8 of the 19 oGPCRs examined can indeed activate G_i/o_ proteins. Specifically, we confirmed previously identified G_i/o_ coupling for the orphan receptors GPR22 (Adams et al., 2008), GPR88 (Dzierba et al., 2015; Jin et al., 2014) and GPRC6A (Pi, Parrill, & Quarles, 2010), even though some reports failed to reproduce G_i/o_ coupling for GPRC6A (Jacobsen et al., 2013). Moreover, we revealed previously unreported robust and significant constitutive activity for GPR156 (8.94 ± 0.40 fold increase over control using the G_s_G_i1_ chimera), GPR137b (4.05 ± 0.45), GPR158 (4.87 ± 0.32), GPR179 (7.97 ± 0.55), and GPRC5D (3.61 ± 0.17) (Figure 4a-b).

The lack of signal obtained transfecting several oGPCRs with any of the tested luciferase reporters may be due to a variety of factors. For example, we demonstrated that ADRA2A receptor was functional in activating wild type or chimeric G proteins (Figure 3b-d), but did not produce a detectable basal G protein signaling (Figure 2d) pointing at a very low level of constitutive activity. Alternatively, the absence of signal could depend on DNA constructs that do not express adequate levels of GPCRs. To test this possibility, we analyzed the expression of our oGPCR library at the protein level by western blot using antibodies directed against C-terminus HA-tag (Figure 4c-d) or myc-tag (Figure 4e). Immunoblots revealed that the expression levels of GPR85 and GPR137 were below detectable threshold, thus providing a possible explanation for their lack of signal (Figure 4c).

### 3.4. Validation of GPR156 constitutive activation of G_i/o_ proteins

Adenylate cyclase represents one of the main intracellular effectors for both G_s_ and G_i_ protein signaling, with G_s_ stimulating cAMP production and G_i_ inhibiting it. To validate the results obtained measuring G_i/o_ constitutive activation by oGPCRs shown in figure 4, we quantified the reduction in cAMP levels induced by treatment with the adenylate cyclase stimulant forskolin in cells overexpressing GPR156, GRM2 or GPRC5B. Using a co-transfected cAMP sensor, we were able to obtain real time measurements of cAMP changes (Figure 5a). As expected, we found that forskolin stimulation of cAMP levels was not affected by overexpression of GPRC5B (107.8 ± 6.5% of CNT; p = 0.622); while overexpression of the positive control GRM2 (54.6 ± 4.7% of CNT) or the orphan receptor GPR156 (65.7 ± 5.2% of CNT) significantly blunted the effect of forskolin (Figure 5a-b). These results confirmed the earlier identified G_i/o_ coupling and high constitutive activity for GPR156, as well as the lack of G_i/o_ signaling for GPRC5B.

**Figure 5.**
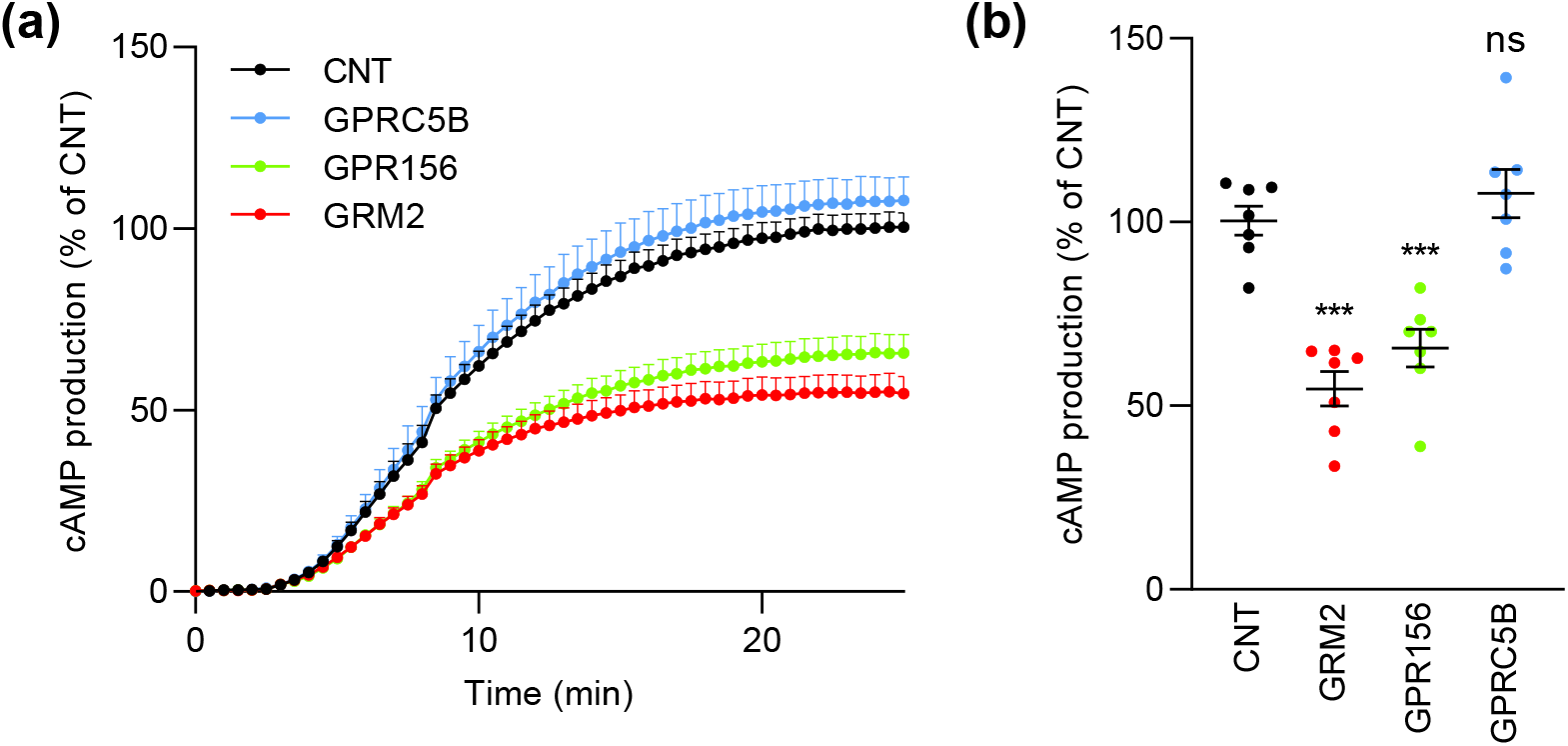
G_i/o_ coupling validation for GPR156. (a) cAMP production induced by 0.5 μM forskolin treatment at 3 minutes. (b) Quantification of the forskolin-induced amplitude reported in panel (a). Cells overexpressing GPR156 and GRM2 show a constitutive inhibition of cAMP production while cells transfected with GPRC5B are not significantly different from control cells transfected with empty vector. Data are shown as means ± SEM (n = 7 independent experiments; one-way ANOVA with Dunnett’s multiple comparisons test, ***p<0.001).

## 4. DISCUSSION

The unique properties of each GPCR together with the plethora of signaling cascades activated makes the development of tailor-made assays a prerequisite for future attempts at profiling oGPCR signaling. Many efforts have been made to create a universal platform for high-throughput screening of GPCR signaling that is independent of G protein coupling (Inoue et al., 2012; Kroeze et al., 2015). For example, the use of quantitative techniques to measure β-arrestin recruitment as a general readout of GPCR activation led to the identification of a number of compounds within a library of 446 molecules acting as agonists or antagonists for class A oGPCRs (Kroeze et al., 2015). However, despite the ability of class C GPCR members to recruit β-arrestins (Iacovelli, Felicioni, Nistico, Nicoletti, & De Blasi, 2014; Mos, Jacobsen, Foster, & Brauner-Osborne, 2019; Mundell, Matharu, Pula, Roberts, & Kelly, 2001; Stoppel et al., 2017), attempts to use this approach to deorphanize this subfamily of oGPCRs were unsuccessful (Kroeze et al., 2015). Similarly, the use of cell-based assays expressing G protein chimeras in G protein knock out cell lines to measure ligand-activated GPCR signaling has recently found a number of applications (Inoue et al., 2019; Okashah et al., 2019). Overall, we expect that a single readout would never be sufficient to detect the activation of every oGPCR without possibly omitting important ligand-receptor pairs. In fact, successful screening efforts will probably need to include multiple alternative readouts. A systematic parallel analysis of GPCR constitutive activity represents a powerful strategy to begin understanding the cell signaling pathways modulated by oGPCRs. Using a novel approach combining luciferase reporter assays with G protein chimeras, here we detected G_i/o_ protein activation by several oGPCRs in absence of ligand stimulation, thereby providing the first evidence for G protein coupling-preference for multiple oGPCRs. This information is crucial in the deorphanization process, as it provides a novel readout in designing platforms to test the activation of oGPCRs allowing for the analysis of libraries of synthetic or endogenous compounds.

In the present study, we did not found evidence of G_s_, G_q_, or G_12/13_ coupling for any of the 19 oGPCRs analyzed, nevertheless, we confirmed G_i/o_ coupling for GPR22, GPR88, and GPRC6A. Strikingly, we observed previously unappreciated G_i/o_ constitutive activities for GPR137b, GPR156, GPR158, GPR179, and GPRC5D. GPR137b expression is restricted to heart, liver, kidney and brain, and it is one of the few GPCRs enriched at lysosomal membranes (Gan et al., 2019; Gao et al., 2012). Proteomics studies of lysosomal membranes also identified several G protein signaling elements including Gαi2, Gβ1, and Gβ2 (Callahan, Bagshaw, & Mahuran, 2009). The functional consequences of activating G_i/o_ signaling responses at the lysosomal membrane remain to be characterized. The group of Pangalos suggested that the class C orphan GPR156 could possibly act as a third GABA_B_ receptor subunit because of their significant sequence homology (Calver et al., 2003). However, functional assays failed to reveal any activation in response to treatments with GABA_B_ receptor agonists in cells expressing GPR156 alone or co-expressing GPR156 with GABA_B1_ or GABA_B2_ receptors (Calver et al., 2003). Searching for alternative ligands, a calcium mobilization assay was used to screen a library of 2500 endogenous GPCR agonists without success (Calver et al., 2003). The extremely high constitutive activity of GPR156 could result in a low dynamic range when performing functional screens and therefore limit the chances to identify possible agonists. At the same time, a high constitutive activity can be a useful tool for the identification of inverse agonists that represent attractive compounds for multiple pharmacotherapies (Berg & Clarke, 2018; Bond & Ijzerman, 2006; Chen et al., 2020). Our screening also revealed G_i/o_ constitutive activation for both GPR158 and GPR179, highly homologous class C receptors. GPR158 is abundantly expressed in several neuronal populations in the brain where it regulates stress-induced depression (Orlandi et al., 2012; Orlandi, Sutton, Muntean, Song, & Martemyanov, 2019; Sutton et al., 2018). While, GPR179 is specifically expressed in the ON-bipolar neurons of the retina (Audo et al., 2012; Orlandi et al., 2012; Peachey et al., 2012). Point mutations in GPR179 gene were identified in patients with congenital stationary night blindness and further animal studies revealed its essential role in night vision (Audo et al., 2012; Peachey et al., 2012; Ray et al., 2014). At the molecular level, both GPR158 and GPR179 has been shown to interact and modulate the activity of a family of R7 Regulator of G protein signaling (R7-RGS) proteins (Orlandi et al., 2012). At the same time GPR179 acts as a scaffold for many components of the post-synaptic mGluR6-G_o_-TRPM1 signaling complex (Orlandi, Cao, & Martemyanov, 2013). Moreover, their long extracellular N-termini have been shown to interact with extracellular matrix components to form trans-synaptic complexes (Condomitti et al., 2018; Dunn, Orlandi, & Martemyanov, 2019; Orlandi et al., 2018). Our results here indicate that GPR158 and GPR179 can simultaneously activate G proteins of the G_i/o_ family while scaffolding R7-RGS proteins, which act as GTPase activating proteins (GAP) for a subset of Gα_i/o_ family members to terminate the G protein signal. These complexes may be required for the timely inactivation of G proteins in response to extracellular events thus limiting the diffusion of activated G proteins to a restricted post-synaptic microenvironment. Alternatively, this receptor complex configuration may limit the diversity of G proteins activated by GPR158 and GPR179 to a subset of Gα_i/o_ family members that are not a suitable substrate for R7-RGS proteins such as Gα_z_. Further studies are needed to investigate the molecular implications of G protein activation by GPR158 in the brain and GPR179 in the retina. The involvement of GPR158 in stress-induced depression makes it an ideal target for development of a novel antidepressants, a desperately needed class of pharmaceuticals. While GPR179 loss of function in congenital stationary night blindness, a debilitating disease without current available treatments, makes it also a relevant candidate for drug discovery ventures. Finally, we detected a previously unreported G_i/o_ coupling for GPRC5D, the last identified member of the class C retinoic acid-inducible receptor family (Brauner-Osborne et al., 2001). GPRC5D is mostly expressed in peripheral tissues and it has recently been associated with cancer (Atamaniuk et al., 2012; Smith et al., 2019). Specifically, its expression was found to be elevated on the surface of malignant cells involved in multiple myeloma, and it represents a viable target for chimeric antigen receptor (CAR) T cell immunotherapy of multiple myeloma (Kodama et al., 2019; Smith et al., 2019). The discovery of signaling pathways triggered by GPRC5D represents therefore an important step forward in the development of therapeutic treatments for white blood plasma cell cancer.

In summary, here we provided a new sensitive strategy to profile constitutive G_i/o_ protein coupling for understudied orphan GPCRs. This approach represents a fundamental advancement in the deorphanization process and will likely accelerate the search for novel GPCR ligands. By screening the entire class C GPCR family, we discovered or confirmed G_i/o_ coupling for 5 out of the 8 orphan members, and, similarly, we revealed 3 G_i/o_ coupled receptors within a subset of class A oGPCRs. Several of these oGPCRs are associated with debilitating neuropsychiatric disorders or they are relevant for treatment of numerous cancers. Hence, improving our understanding of the biology of such receptors has clinical relevance and it is essential in the drug discovery process.

## Acknowledgments

For providing essential cDNA constructs, we are sincerely grateful for Dr. Kirill A. Martemyanov (The Scripps Research Institute, Jupiter, FL), Dr. Paul J. Kammermeier (University of Rochester, NY), Dr. Nevin A. Lambert (Augusta University, GA), Dr. Mark H. Ginsberg (University of California San Diego, CA), Dr. Bryan L. Roth (University of North Carolina, NC), Dr. Bruce R. Conklin (University of California San Francisco, CA), Dr. Robert J. Lucas (University of Manchester, UK), and Dr. Erik Procko (University of Illinois at Urbana, IL). We also would like to thank Dr. Henry A. Dunn (The Scripps Research Institute, Jupiter, FL) for comments and fruitful discussions.

## Author Contributions

Experimental investigation and data analysis, L.R.W. and C.O; Conceptualization, C.O.; writing and editing—original draft preparation, C.O.; All authors have read and agreed to the published version of the manuscript.

## References

Adams, J. W., Wang, J., Davis, J. R., Liaw, C., Gaidarov, I., Gatlin, J., … Connolly, D. (2008). Myocardial expression, signaling, and function of GPR22: a protective role for an orphan G protein-coupled receptor. Am J Physiol Heart Circ Physiol, 295(2), H509–521. doi:10.1152/ajpheart.00368.2008

Ahmad, R., Wojciech, S., & Jockers, R. (2015). Hunting for the function of orphan GPCRs - beyond the search for the endogenous ligand. Br J Pharmacol, 172(13), 3212–3228. doi:10.1111/bph.12942

Atamaniuk, J., Gleiss, A., Porpaczy, E., Kainz, B., Grunt, T. W., Raderer, M., … Gaiger, A. (2012). Overexpression of G protein-coupled receptor 5D in the bone marrow is associated with poor prognosis in patients with multiple myeloma. Eur J Clin Invest, 42(9), 953–960. doi:10.1111/j.1365-2362.2012.02679.x

Audo, I., Bujakowska, K., Orhan, E., Poloschek, C. M., Defoort-Dhellemmes, S., Drumare, I., … Zeitz, C. (2012). Whole-exome sequencing identifies mutations in GPR179 leading to autosomal-recessive complete congenital stationary night blindness. Am J Hum Genet, 90(2), 321–330. doi:10.1016/j.ajhg.2011.12.007

Ballister, E. R., Rodgers, J., Martial, F., & Lucas, R. J. (2018). A live cell assay of GPCR coupling allows identification of optogenetic tools for controlling Go and Gi signaling. BMC Biol, 16(1), 10. doi:10.1186/s12915-017-0475-2

Berg, K. A., & Clarke, W. P. (2018). Making Sense of Pharmacology: Inverse Agonism and Functional Selectivity. Int J Neuropsychopharmacol, 21(10), 962–977. doi:10.1093/ijnp/pyy071

Bond, R. A., & Ijzerman, A. P. (2006). Recent developments in constitutive receptor activity and inverse agonism, and their potential for GPCR drug discovery. Trends Pharmacol Sci, 27(2), 92–96. doi:10.1016/j.tips.2005.12.007

Brauner-Osborne, H., Jensen, A. A., Sheppard, P. O., Brodin, B., Krogsgaard-Larsen, P., & O’Hara, P. (2001). Cloning and characterization of a human orphan family C G-protein coupled receptor GPRC5D. Biochim Biophys Acta, 1518(3), 237–248. doi:10.1016/s0167-4781(01)00197-x

Callahan, J. W., Bagshaw, R. D., & Mahuran, D. J. (2009). The integral membrane of lysosomes: its proteins and their roles in disease. J Proteomics, 72(1), 23–33. doi:10.1016/j.jprot.2008.11.007

Calver, A. R., Michalovich, D., Testa, T. T., Robbins, M. J., Jaillard, C., Hill, J., … Pangalos, M. N. (2003). Molecular cloning and characterisation of a novel GABAB-related G-protein coupled receptor. Brain Res Mol Brain Res, 110(2), 305–317. doi:10.1016/s0169-328x(02)00662-9

Chen, Z., Chen, H., Zhang, Z., Ding, P., Yan, X., Li, Y., … Xu, J. (2020). Discovery of novel liver X receptor inverse agonists as lipogenesis inhibitors. Eur J Med Chem, 206, 112793. doi:10.1016/j.ejmech.2020.112793

Cheng, Z., Garvin, D., Paguio, A., Stecha, P., Wood, K., & Fan, F. (2010). Luciferase Reporter Assay System for Deciphering GPCR Pathways. Curr Chem Genomics, 4, 84–91. doi:10.2174/1875397301004010084

Condomitti, G., Wierda, K. D., Schroeder, A., Rubio, S. E., Vennekens, K. M., Orlandi, C., … de Wit, J. (2018). An Input-Specific Orphan Receptor GPR158-HSPG Interaction Organizes Hippocampal Mossy Fiber-CA3 Synapses. Neuron, 100(1), 201–215 e209. doi:10.1016/j.neuron.2018.08.038

Conklin, B. R., Farfel, Z., Lustig, K. D., Julius, D., & Bourne, H. R. (1993). Substitution of three amino acids switches receptor specificity of Gq alpha to that of Gi alpha. Nature, 363(6426), 274–276. doi:10.1038/363274a0

Corder, G., Doolen, S., Donahue, R. R., Winter, M. K., Jutras, B. L., He, Y., … Taylor, B. K. (2013). Constitutive mu-opioid receptor activity leads to long-term endogenous analgesia and dependence. Science, 341(6152), 1394–1399. doi:10.1126/science.1239403

Coward, P., Chan, S. D., Wada, H. G., Humphries, G. M., & Conklin, B. R. (1999). Chimeric G proteins allow a high-throughput signaling assay of Gi-coupled receptors. Anal Biochem, 270(2), 242–248. doi:10.1006/abio.1999.4061

Damian, M., Marie, J., Leyris, J. P., Fehrentz, J. A., Verdie, P., Martinez, J., … Mary, S. (2012). High constitutive activity is an intrinsic feature of ghrelin receptor protein: a study with a functional monomeric GHS-R1a receptor reconstituted in lipid discs. J Biol Chem, 287(6), 3630–3641. doi:10.1074/jbc.M111.288324

Doornbos, M. L. J., Van der Linden, I., Vereyken, L., Tresadern, G., Ap, I. J., Lavreysen, H., & Heitman, L. H. (2018). Constitutive activity of the metabotropic glutamate receptor 2 explored with a whole-cell label-free biosensor. Biochem Pharmacol, 152, 201–210. doi:10.1016/j.bcp.2018.03.026

Dunn, H. A., Orlandi, C., & Martemyanov, K. A. (2019). Beyond the Ligand: Extracellular and Transcellular G Protein-Coupled Receptor Complexes in Physiology and Pharmacology. Pharmacol Rev, 71(4), 503–519. doi:10.1124/pr.119.018044

Dzierba, C. D., Bi, Y., Dasgupta, B., Hartz, R. A., Ahuja, V., Cianchetta, G., … Macor, J. E. (2015). Design, synthesis, and evaluation of phenylglycinols and phenyl amines as agonists of GPR88. Bioorg Med Chem Lett, 25(7), 1448–1452. doi:10.1016/j.bmcl.2015.01.036

Flock, T., Hauser, A. S., Lund, N., Gloriam, D. E., Balaji, S., & Babu, M. M. (2017). Selectivity determinants of GPCR-G-protein binding. Nature, 545(7654), 317–322. doi:10.1038/nature22070

Gan, L., Seki, A., Shen, K., Iyer, H., Han, K., Hayer, A., … Meyer, T. (2019). The lysosomal GPCR-like protein GPR137B regulates Rag and mTORC1 localization and activity. Nat Cell Biol, 21(5), 614–626. doi:10.1038/s41556-019-0321-6

Gao, J., Xia, L., Lu, M., Zhang, B., Chen, Y., Xu, R., & Wang, L. (2012). TM7SF1 (GPR137B): a novel lysosome integral membrane protein. Mol Biol Rep, 39(9), 8883–8889. doi:10.1007/s11033-012-1755-0

Hauser, A. S., Attwood, M. M., Rask-Andersen, M., Schioth, H. B., & Gloriam, D. E. (2017). Trends in GPCR drug discovery: new agents, targets and indications. Nat Rev Drug Discov, 16(12), 829–842. doi:10.1038/nrd.2017.178

Hollins, B., Kuravi, S., Digby, G. J., & Lambert, N. A. (2009). The c-terminus of GRK3 indicates rapid dissociation of G protein heterotrimers. Cell Signal, 21(6), 1015–1021. doi:10.1016/j.cellsig.2009.02.017

Iacovelli, L., Felicioni, M., Nistico, R., Nicoletti, F., & De Blasi, A. (2014). Selective regulation of recombinantly expressed mGlu7 metabotropic glutamate receptors by G protein-coupled receptor kinases and arrestins. Neuropharmacology, 77, 303–312. doi:10.1016/j.neuropharm.2013.10.013

Inoue, A., Ishiguro, J., Kitamura, H., Arima, N., Okutani, M., Shuto, A., … Aoki, J. (2012). TGFalpha shedding assay: an accurate and versatile method for detecting GPCR activation. Nat Methods, 9(10), 1021–1029. doi:10.1038/nmeth.2172

Inoue, A., Raimondi, F., Kadji, F. M. N., Singh, G., Kishi, T., Uwamizu, A., … Russell, R. B. (2019). Illuminating G-Protein-Coupling Selectivity of GPCRs. Cell, 177(7), 1933–1947 e1925. doi:10.1016/j.cell.2019.04.044

Jacobsen, S. E., Norskov-Lauritsen, L., Thomsen, A. R., Smajilovic, S., Wellendorph, P., Larsson, N. H., … Brauner-Osborne, H. (2013). Delineation of the GPRC6A receptor signaling pathways using a mammalian cell line stably expressing the receptor. J Pharmacol Exp Ther, 347(2), 298–309. doi:10.1124/jpet.113.206276

Jin, C., Decker, A. M., Huang, X. P., Gilmour, B. P., Blough, B. E., Roth, B. L., … Zhang, X. P. (2014). Synthesis, pharmacological characterization, and structure-activity relationship studies of small molecular agonists for the orphan GPR88 receptor. ACS Chem Neurosci, 5(7), 576–587. doi:10.1021/cn500082p

Kodama, T., Kochi, Y., Nakai, W., Mizuno, H., Baba, T., Habu, K., … Akashi, K. (2019). Anti-GPRC5D/CD3 Bispecific T-Cell-Redirecting Antibody for the Treatment of Multiple Myeloma. Mol Cancer Ther, 18(9), 1555–1564. doi:10.1158/1535-7163.MCT-18-1216

Kroeze, W. K., Sassano, M. F., Huang, X. P., Lansu, K., McCorvy, J. D., Giguere, P. M., … Roth, B. L. (2015). PRESTO-Tango as an open-source resource for interrogation of the druggable human GPCRome. Nat Struct Mol Biol, 22(5), 362–369. doi:10.1038/nsmb.3014

Masuho, I., Martemyanov, K. A., & Lambert, N. A. (2015). Monitoring G Protein Activation in Cells with BRET. Methods Mol Biol, 1335, 107–113. doi:10.1007/978-1-4939-2914-6_8

Mos, I., Jacobsen, S. E., Foster, S. R., & Brauner-Osborne, H. (2019). Calcium-Sensing Receptor Internalization Is beta-Arrestin-Dependent and Modulated by Allosteric Ligands. Mol Pharmacol, 96(4), 463–474. doi:10.1124/mol.119.116772

Mundell, S. J., Matharu, A. L., Pula, G., Roberts, P. J., & Kelly, E. (2001). Agonist-induced internalization of the metabotropic glutamate receptor 1a is arrestin- and dynamin-dependent. J Neurochem, 78(3), 546–551. doi:10.1046/j.1471-4159.2001.00421.x

Ngo, T., Coleman, J. L., & Smith, N. J. (2015). Using constitutive activity to define appropriate high-throughput screening assays for orphan g protein-coupled receptors. Methods Mol Biol, 1272, 91–106. doi:10.1007/978-1-4939-2336-6_7

Okashah, N., Wan, Q., Ghosh, S., Sandhu, M., Inoue, A., Vaidehi, N., & Lambert, N. A. (2019). Variable G protein determinants of GPCR coupling selectivity. Proc Natl Acad Sci U S A, 116(24), 12054–12059. doi:10.1073/pnas.1905993116

Orlandi, C., Cao, Y., & Martemyanov, K. A. (2013). Orphan receptor GPR179 forms macromolecular complexes with components of metabotropic signaling cascade in retina ON-bipolar neurons. Invest Ophthalmol Vis Sci, 54(10), 7153–7161. doi:10.1167/iovs.13-12907

Orlandi, C., Omori, Y., Wang, Y., Cao, Y., Ueno, A., Roux, M. J., … Martemyanov, K. A. (2018). Transsynaptic Binding of Orphan Receptor GPR179 to Dystroglycan-Pikachurin Complex Is Essential for the Synaptic Organization of Photoreceptors. Cell Rep, 25(1), 130–145 e135. doi:10.1016/j.celrep.2018.08.068

Orlandi, C., Posokhova, E., Masuho, I., Ray, T. A., Hasan, N., Gregg, R. G., & Martemyanov, K. A. (2012). GPR158/179 regulate G protein signaling by controlling localization and activity of the RGS7 complexes. J Cell Biol, 197(6), 711–719. doi:10.1083/jcb.201202123

Orlandi, C., Sutton, L. P., Muntean, B. S., Song, C., & Martemyanov, K. A. (2019). Homeostatic cAMP regulation by the RGS7 complex controls depression-related behaviors. Neuropsychopharmacology, 44(3), 642–653. doi:10.1038/s41386-018-0238-y

Pandy-Szekeres, G., Munk, C., Tsonkov, T. M., Mordalski, S., Harpsoe, K., Hauser, A. S., … Gloriam, D. E. (2018). GPCRdb in 2018: adding GPCR structure models and ligands. Nucleic Acids Res, 46(D1), D440–D446. doi:10.1093/nar/gkx1109

Park, J., Selvam, B., Sanematsu, K., Shigemura, N., Shukla, D., & Procko, E. (2019). Structural architecture of a dimeric class C GPCR based on co-trafficking of sweet taste receptor subunits. J Biol Chem, 294(13), 4759–4774. doi:10.1074/jbc.RA118.006173

Peachey, N. S., Ray, T. A., Florijn, R., Rowe, L. B., Sjoerdsma, T., Contreras-Alcantara, S., … Gregg, R. G. (2012). GPR179 is required for depolarizing bipolar cell function and is mutated in autosomal-recessive complete congenital stationary night blindness. Am J Hum Genet, 90(2), 331–339. doi:10.1016/j.ajhg.2011.12.006

Pi, M., Parrill, A. L., & Quarles, L. D. (2010). GPRC6A mediates the non-genomic effects of steroids. J Biol Chem, 285(51), 39953–39964. doi:10.1074/jbc.M110.158063

Ray, T. A., Heath, K. M., Hasan, N., Noel, J. M., Samuels, I. S., Martemyanov, K. A., … Gregg, R. G. (2014). GPR179 is required for high sensitivity of the mGluR6 signaling cascade in depolarizing bipolar cells. J Neurosci, 34(18), 6334–6343. doi:10.1523/JNEUROSCI.4044-13.2014

Rosenbaum, D. M., Rasmussen, S. G., & Kobilka, B. K. (2009). The structure and function of G-protein-coupled receptors. Nature, 459(7245), 356–363. doi:10.1038/nature08144

Smith, E. L., Harrington, K., Staehr, M., Masakayan, R., Jones, J., Long, T. J., … Brentjens, R. J. (2019). GPRC5D is a target for the immunotherapy of multiple myeloma with rationally designed CAR T cells. Sci Transl Med, 11(485). doi:10.1126/scitranslmed.aau7746

Sriram, K., & Insel, P. A. (2018). G Protein-Coupled Receptors as Targets for Approved Drugs: How Many Targets and How Many Drugs? Mol Pharmacol, 93(4), 251–258. doi:10.1124/mol.117.111062

Stoppel, L. J., Auerbach, B. D., Senter, R. K., Preza, A. R., Lefkowitz, R. J., & Bear, M. F. (2017). beta-Arrestin2 Couples Metabotropic Glutamate Receptor 5 to Neuronal Protein Synthesis and Is a Potential Target to Treat Fragile X. Cell Rep, 18(12), 2807–2814. doi:10.1016/j.celrep.2017.02.075

Sutton, L. P., Orlandi, C., Song, C., Oh, W. C., Muntean, B. S., Xie, K., … Martemyanov, K. A. (2018). Orphan receptor GPR158 controls stress-induced depression. Elife, 7. doi:10.7554/eLife.33273

Wang, D., Stoveken, H. M., Zucca, S., Dao, M., Orlandi, C., Song, C., … Martemyanov, K. A. (2019). Genetic behavioral screen identifies an orphan anti-opioid system. Science, 365(6459), 1267–1273. doi:10.1126/science.aau2078

Watkins, L. R., & Orlandi, C. (2020). Orphan G Protein Coupled Receptors in Affective Disorders. Genes (Basel), 11(6). doi:10.3390/genes11060694

